# Auricular transcutaneous vagus nerve stimulation acutely modulates brain connectivity in mice

**DOI:** 10.1101/2021.11.15.468252

**Authors:** Cecilia Brambilla-Pisoni, Emma Muñoz-Moreno, Ianire Gallego-Amaro, Rafael Maldonado, Antoni Ivorra, Guadalupe Soria, Andrés Ozaita

## Abstract

**Background:** Brain electrical stimulation techniques take advantage of the intrinsic plasticity of the nervous system, opening a wide range of therapeutic applications. Vagus nerve stimulation (VNS) is an approved adjuvant for drug-resistant epilepsy and depression. Its non-invasive form, auricular transcutaneous VNS (atVNS), is under investigation for applications, including cognitive improvement.

**Objective:** We aimed to study the effects of atVNS on brain connectivity, under conditions that improved memory persistence in CD-1 male mice.

**Methods:** Acute atVNS in the *cymba conchae* of the left ear was performed using a standard stimulation protocol under light isoflurane anesthesia, immediately or 3 h after the training/familiarization phase of the novel object-recognition memory test (NORT). Another cohort of mice was used for bilateral c-Fos analysis after atVNS administration. Spearman correlation of c-Fos density between each pair of the thirty brain regions analyzed allowed obtaining the network of significant functional connections in stimulated and non-stimulated control brains.

**Results:** NORT performance was enhanced when atVNS was delivered just after, but not 3 h after, the familiarization phase of the task. No alterations in c-Fos density were associated to electrostimulation, but a significant effect of atVNS was observed on c-Fos-based functional connectivity. atVNS induced a clear reorganization of the network, increasing the inter-hemisphere connections and the connectivity of locus coeruleus.

**Conclusion:** Our results provide new insights in the effects of atVNS on memory performance and brain connectivity extending our knowledge of the biological mechanisms of bioelectronics in medicine.

**Highlights:**

- atVNS, delivered immediately after NORT training phase, improves memory persistence
- atVNS did not promote significant changes in brain c-Fos density
- atVNS induced a significant reorganization of c-Fos-based functional brain network
- atVNS produced an enhancement in correlated activity between hemispheres
- atVNS did not engage the prefrontal-retrosplenial axis, characteristic of the DMN

## Introduction

Nowadays, brain stimulation devices have gained significant interest in the scientific community and have received European Medicine Agency (EMA) and US Food and Drug Administration (FDA) approvals for different therapeutic purposes. An approach to achieve brain stimulation is through vagus nerve afferents. Indeed, vagus nerve stimulation (VNS) has become an interesting strategy to help handle drug-resistant epilepsy and depression [1]. In this context, transcutaneous VNS (tVNS), given its non-invasiveness, has received substantial attention. tVNS can be applied to different locations, such as in the neck [2] or in the *cymba conchae* of the external ear [3], and, similar to the invasive form of VNS, has already been approved as an adjuvant in clinical applications, such as for the treatment of drug-resistant epilepsy and depression [4,5]. Surprisingly, the direct effect of tVNS over brain function is far from understood. Renewed attention to this electrostimulation technique derives from its neuromodulatory effect of cognitive processes. The vagus nerve afferent fibers in the brainstem end in the nucleus of the solitary tract (NTS) [6], a relevant relay area for visceral information. From there, afferent information is distributed to many brain regions, including the *locus coeruleus* (LC) [7]. The LC provides a widespread innervation to the hippocampus, amygdala and prefrontal cortex, among other regions [6]. In this regard, it has been postulated that the LC could regulate cognition through the release of norepinephrine and dopamine in memory processing areas [6,8], mimicking those physiological processes involved in attention-driven cognition [9-11], but whether such interaction results in the modulation of brain networks is not completely understood. Among those brain networks supporting brain activity, the default mode network (DMN), a brain network predominantly active when the brain is not engaged in an attention-driven task [12,13], has received much consideration due to the reproducibility in its detection in clinical and preclinical settings [13]. The DMN is therefore disengaged during periods of active brain function, as during attention-associated periods, and it has been described to be modulated by tVNS in patients with mild or moderate depressive symptoms [14]. Focusing on the cognitive functions associated to memory, our group already reported that auricular tVNS (atVNS) enhanced novel object-recognition (NOR) memory persistence in naïve CD-1 mice [15]. The brain mechanisms recruited by atVNS to enhance memory performance have not been described, so we further explored the cellular outcome of atVNS under conditions that potentiate object-recognition memory persistence. Firstly, we assessed whether a critical time window for atVNS efficacy may limit its effect on favoring object-recognition memory. Secondly, we analyzed the expression of the immediate early gene c-Fos as an approach to study brain activity in discrete brain regions [16]. In line with the current view proposing that specific cognitive functions are supported by a network of functionally connected brain regions, rather than isolated areas, together with the region-specific analysis of c-Fos, we also evaluated the c-Fos-based functional network [17]. We focused our analysis in areas of the brainstem, hippocampus, amygdala, thalamus and frontal/dorsal cortex, some of which are components of the DMN. Starting from these c-Fos data, we analyzed and estimated the functional connectivity network based on c-Fos density for both stimulated and non-stimulated brains to find significant changes in network connectivity patterns in atVNS condition.

## Materials and methods

### Animals

Young-adult male CD-1 mice (10 weeks old) were purchased from Charles River Laboratories (France). All experimental mice were bred at the Barcelona Biomedical Research Park (PRBB) Animal Facility. All animal procedures were conducted in accordance with the standard ethical guidelines (European Communities Directive 2010/63/EU). Mice were housed in a temperature-controlled (21 ± 1 °C) and humidity-controlled (55 ± 10%) environment. Lighting was maintained at 12 hours’ cycles (on at 8 AM and off at 8 PM). Food and water were available *ad libitum*. Mice were handled for 1 week before starting the experiment and were randomly distributed among experimental groups. All procedures were performed by experimenters blind to the experimental conditions.

### Behavioral and electrostimulation procedures

The electrostimulation was performed at the time points indicated below after the familiarization/training phase in the novel object-recognition test (NORT), following a similar approach to that described previously [15]. Briefly, on the habituation phase performed in day 1, mice were habituated to an empty V-shape maze (V-maze) for 9 min. The next day, on the training phase performed in day 2, mice were presented in the V-maze to two identical objects for 9 min, each object at the end of the maze corridors. Immediately after the familiarization phase (atVNS (0h), n=11) or 3 h after the familiarization phase (atVNS (3h), n=12) mice were anesthetized with isoflurane (1.5%) in 0.8 L/min O_2_ during 30 min, and subjected or not to atVNS. Normothermic conditions were maintained during anesthesia with a heating pad. For atVNS (0h) and atVNS (3h) conditions, a bipolar electrode (described in 15) was placed in the *cymbae concha* of the left ear. Rectangular biphasic pulses were delivered with a Beurer EM49 stimulator (Beurer, Germany). The stimulation parameters were: 1 mA, 20 Hz, 30 s ON and 5 min OFF, total length of 30 min, with a 320 μs pulse width. For the No stimulation condition (n=12), mice were anesthetized for 30 min immediately after the NORT familiarization, but no electrical stimulation was delivered. 48 h after the NORT familiarization phase and the atVNS or No stimulation procedures, mice were tested for 9 min in the V-maze, substituting one of the familiar object for a new one, to assess memory performance.

### Tissue preparation for immunofluorescence

In another batch of animals, the exact same NORT+atVNS or NORT+No stimulation protocols described above were followed, returning the mice to their home-cage afterwards. Ninety min following the completion of the NORT+atVNS (similar to atVNS (0h) condition, n=8) or the NORT+No stimulation (No stimulation condition, n=8), mice were deeply anesthetized by intraperitoneal injection (0.2 mL/10 g of body weight) of a mixture of ketamine (100 mg/kg) and xylazine (20 mg/kg) prior to intracardiac perfusion with 4% paraformaldehyde in 0.1M Na_2_HPO_4_/0.1M NaH_2_PO_4_ buffer (PB), pH 7.5, delivered with a peristaltic pump at 19 mL/min flow for 3 min. Subsequently, brain was extracted and post-fixed in the same fixative solution for 24 h and transferred to a solution of 30% sucrose in PB overnight at 4ºC. After post-fixation the brains were marked in the right hemisphere to preserve laterality in the subsequent measures. Coronal sections of 30 μm were obtained on a freezing microtome and stored in a solution of 5% sucrose at 4ºC until used.

### Immunofluorescence

Sections from the No stimulation and atVNS groups were processed in parallel for immunofluorescence. Briefly, free-floating brain slices were rinsed in PB, blocked in a solution containing 3% normal goat serum (GS) (S-1000-20, Vector Laboratories, Inc., CA, USA) and 0.3% Triton X-100 (T) in PB (GS-T-PB) at room temperature for 2 h, and incubated overnight in the same solution with the primary antibody to c-Fos (sc-7202, 1:1000, rabbit, Santa Cruz Biotechnology) and, only for LC slices, with tyrosine hydroxylase (T1299, 1:1,000, mouse, Sigma-Aldrich) at 4ºC. The next day, after 3 rinses in PB, sections were incubated at room temperature with the secondary antibody AlexaFluor-555 goat anti-rabbit (ab150078, 1:1,000, Abcam) and, only for LC slices, with AlexaFluor-488 goat anti-mouse (115-545-003, 1:1,000, Jackson ImmunoResearch Laboratories Inc.) for 2 h. After incubation, sections were rinsed and mounted immediately after onto glass slides coated with gelatin in Fluoromont-G with 4’,6-diamidino-2-phenylindole (DAPI) (00-4959-52, Invitrogen, Thermo Fisher Scientific, MA, USA) as counterstaining.

### c-Fos quantification

c-Fos density was analyzed in thirty brain regions (fifteen per hemisphere), taking into account brain laterality. Analyzed brain regions included (from frontal to caudal): cingulate cortex (Cg), prelimbic cortex (PrL), infralimbic cortex (IL) (coordinates relative to Bregma: 1.94 mm to 1.54 mm), dentate gyrus (DG), CA1 and CA3 areas of the hippocampus (from Bregma: –1.46 mm to –1.82 mm), basolateral amygdala (BLA), lateral amygdala (LA) and central amygdala (Cea) (from Bregma: –1.46 mm to –1.82 mm), paraventricular nucleus of the thalamus (from Bregma: –1-46 mm to –1.82 mm), anterior and posterior retrosplenial cortex (RSP, pRSP) (from Bregma: -1.46 mm to -2.92 mm), *locus coeruleus* (LC) (from Bregma: –5.34 mm to –5.68 mm), nucleus of the solitary tract (NTS) and dorsal vagal nucleus (DMX) (from Bregma: –7.32 mm to –7.64) (Supplementary Fig. 1). Immunostained brain sections were analyzed with a ×10 objective using a Leica DMR microscope (DM6000B, Leica Microsystems, Wetzlar, Germany) equipped with a digital camera Leica DFX 3000FX (Leica Microsystems). The borders of all regions were defined manually according to the mouse brain atlas [18]. For prelimbic, infralimbic and cingulate cortexes analysis, a 430-μm-sided square region of interest (ROI) was delimited for quantification. For amygdala and dorsal hippocampus analysis, the DAPI signal was used for the delimitation of the areas in each image for quantification. For the LC, the tyrosine hydroxylase signal was used for the delimitation of the area in each image for quantification. The images were processed using ImageJ software [19]. c-Fos-positive cells in each brain areas were quantified manually using the cell counter plugin of ImageJ software. The average number of c-Fos-positive cells on four determinations for each brain area on each hemisphere was calculated for each mouse. The c-Fos density for each region was quantified dividing the number of c-Fos-positive cells to the area considered for each region (c-Fos+/mm^2^) (see Supplementary Fig. 1 for representative examples).

### Generation of functional connectivity network

Functional network was estimated for each condition based on correlation between regional c-Fos density, considering that a functional connection exists between two regions if their activity co-varies [17,20]. Therefore, within each experimental group (No stimulation and atVNS), pair-wise Spearman correlation coefficient between each pair of regional c-Fos density were computed. In this way, a correlation matrix was obtained from each condition representing the correlation coefficients between all thirty brain regions analyzed, taking into account brain laterality. By considering only significant correlations (p<0.05), both positive and negative, we obtained the task-related functional network for atVNS and No stimulation conditions. These networks were represented by circos plots, using a custom R-code (R version 4.0.4) [21]. Finally, we computed the z-Fisher transform of significant positive correlation coefficients, as a measure of connectivity strength between nodes (z-score), for both conditions and displayed them by Kamada-Kawai graphs using NetworkX graph python package (NetworkX version 2.5.1) [22] to visualize network organization.

### Network analysis

First, the total functional connectivity strengths for all the possible connections were compared between atVNS and No stimulation conditions, considering z-score. Likewise, for LC region we compared its connectivity strength with all the other evaluated regions between atVNS and non-stimulated networks.

To have a global characterization of the task-related functional connectivity we also computed graph metrics on the network of significant positive connectivity strengths using Brain Connectivity Toolbox (BCT) [23]. In particular, global efficiency, average clustering, average strength and average degree of the network were estimated. In addition, regional network metrics such as nodal strength and nodal degree coefficients were also computed. To compare network organization and the relevance of each region in the functional network, regional metrics were normalized to the maximum in the network and ordered from higher to lower value to identify network hubs [24].

### Statistical analysis

Data were analyzed with STATISTICA (StatSoft) software using one-way analysis of variance (ANOVA) for multiple comparison of parametric variables. Kruskal-Wallis test was used for non-parametric variables. Subsequent post-hoc analysis (Newman-Keuls) was used when required to reveal significant interaction between factors. The artwork was designed using GraphPad Prism 7. Comparisons were considered statistically significant when *p* < 0.05. Data are represented as mean ± s.e.m.

## Results

### Object-recognition memory enhancement by acute atVNS depends on the time of administration

atVNS was administered after the familiarization phase of the NORT at two different time points, immediately after (atVNS (0h) group) or 3 h after (atVNS (3h) group). As control we intermingled another batch of animals that was similarly handled but did not receive the electrostimulation procedure (No stimulation group). We found that an acute session of atVNS, delivered immediately after the NORT familiarization phase, significantly improved object-recognition memory performance at 48 h (Fig. 1A) compared to atVNS (3h) and No stimulation conditions which showed similar results with no enhancement in memory persistence. This result indicates that the modulation of object-recognition memory persistence depends on the time of application after familiarization/training, with a critical time window for effective action of atVNS.

**Figure 1.**
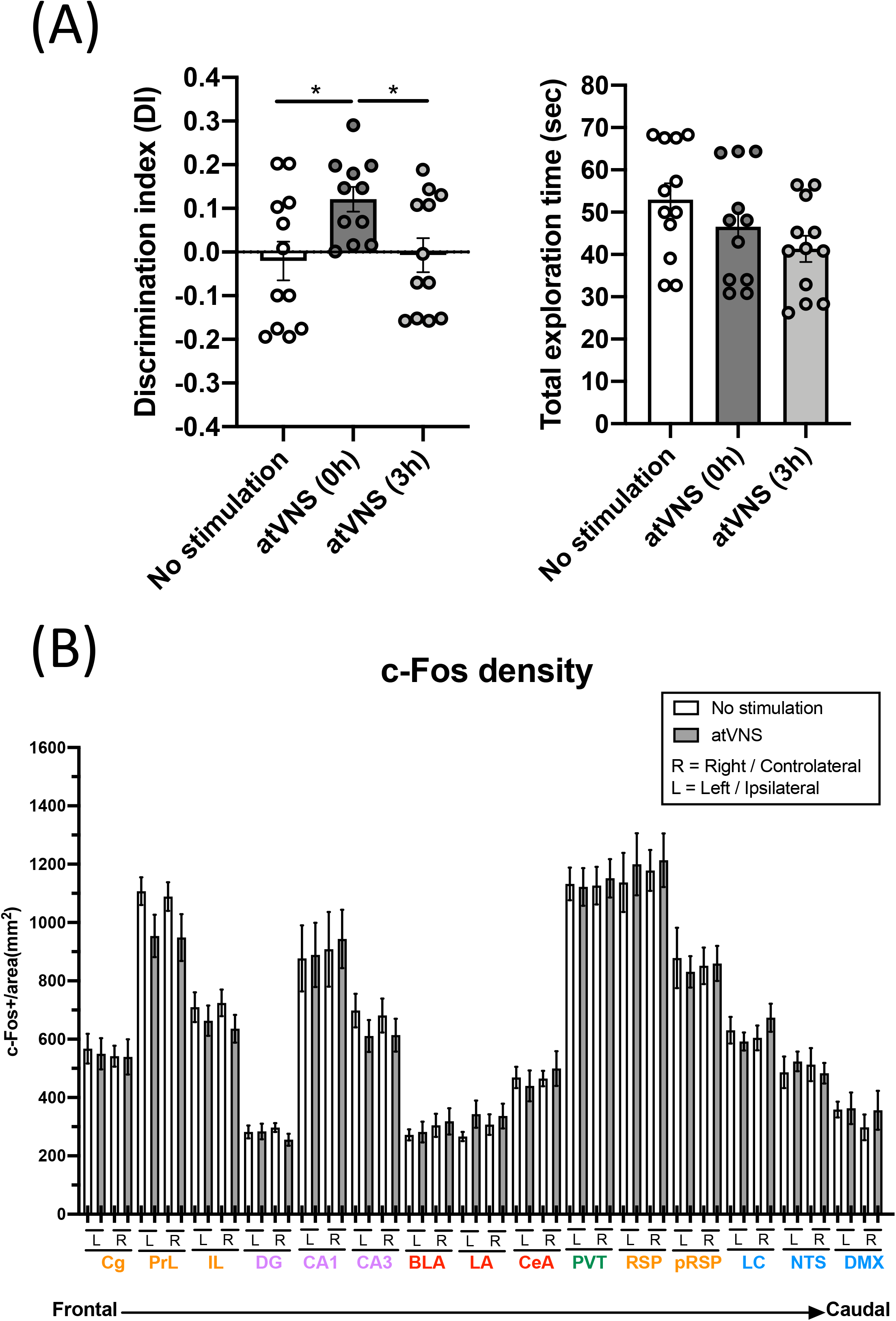
**(A)** atVNS improves object-recognition memory persistence in naïve mice when administered immediately after the familiarization phase of the novel object-recognition test (NORT). Discrimination index and total exploration time in NORT for atVNS (0h), atVNS (3h) and No stimulation conditions in naïve CD-1 mice (atVNS (0h) condition, n=11; atVNS (3h) condition, n=12; No stimulation condition, n=12). *p<0.05 by one-way ANOVA. **(B)** c-Fos density in No stimulation and atVNS conditions, separating contralateral (right, R) and ipsilateral (left, L) sides according to the site of the stimulation. The brain regions analyzed are organized from frontal to caudal and grouped in cortical (orange), hippocampal (purple), amygdalar (red), thalamic (green) and brainstem (blue) groups.

### c-Fos density is not significantly modified in various brain regions after atVNS

Brain samples from NORT+No stimulation and NORT+atVNS (0h) conditions were obtained 90 min after sham or atVNS handling to match the peak of expression of c-Fos. We focused the analysis of c-Fos density in areas involved in novel object-recognition memory processing. Laterality was considered in the analysis.

Notably, c-Fos density analysis did not reveal significant differences between experimental conditions in any of the areas considered, although prelimbic and infralimbic regions and CA3 area showed a non-significant trend to reduce the density of c-Fos positive cells after atVNS (Fig. 1B). This result showed that atVNS is not associated to localized regional changes, and therefore we investigated whether its effects are related to a network reorganization of brain functioning.

### Inferred brain connectivity is relevantly modulated by atVNS

To gain deeper insights into the functional connections within the set of brain regions in our analysis, we computed the connectivity matrices for each experimental group (Supplementary Fig. 2). Comparing both connectivity matrices, an overall effect of atVNS on the relation of c-Fos density among different brain areas was observed, with a higher percentage of positive correlations under atVNS condition. When we statistically compared the strengths of all the connections between both conditions we observed a significant increase in the total connectivity (No stimulation: 0.11 ± 0.029; atVNS: 0.48 ± 0.022; t(868) = 10.05; p < 0.0001) induced by atVNS (Fig 2).

**Figure 2.**
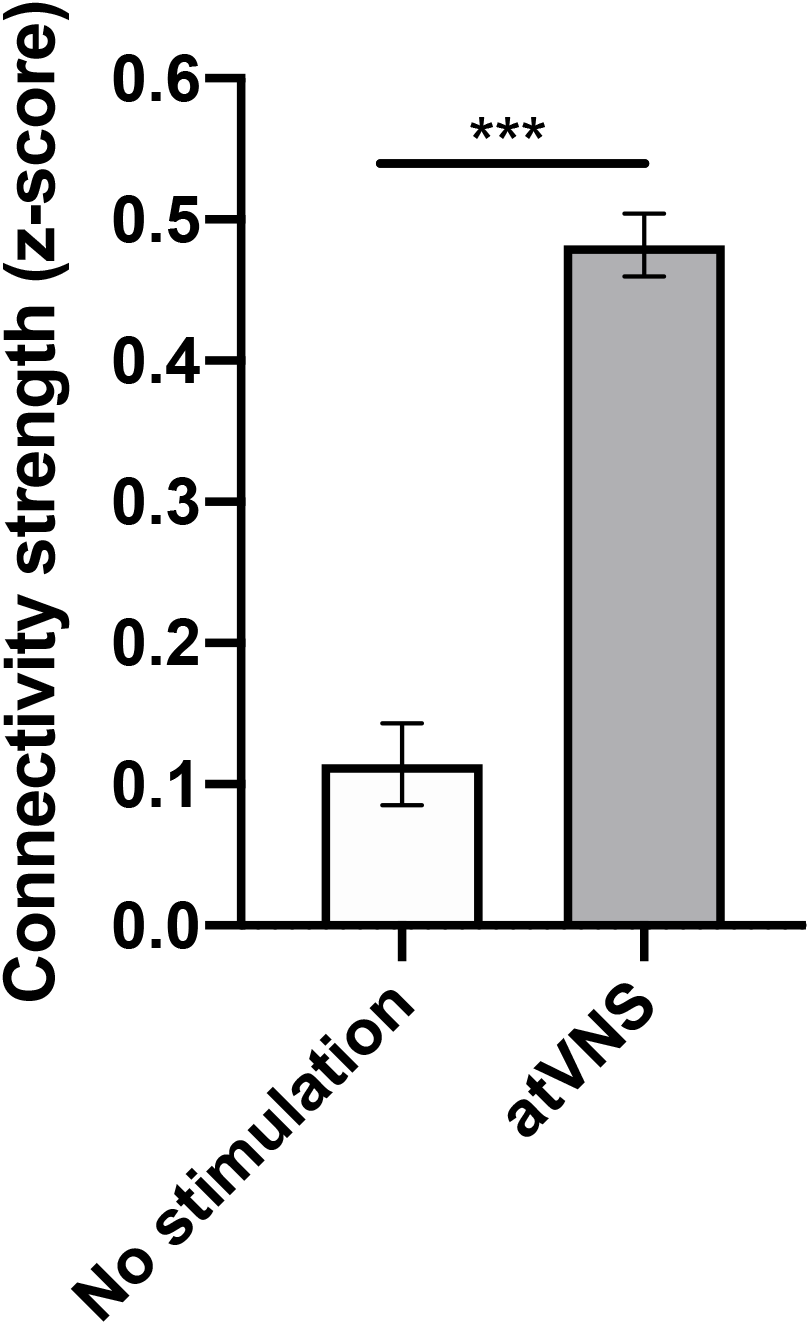
Effects of atVNS on global network connectivity. Total connectivity presented as z-score, comparing No stimulation and atVNS conditions. *** p<0.001 by Kruskal-Wallis test.

Subsequently, we focused on the networks showing significant correlations (p-value < 0.05), that are represented in circos plot of Fig. 3A for each condition. This analysis revealed that atVNS coupled the activity of left and right LC, also increasing its correlated activity with the dentate gyrus. Furthermore, the electrostimulation reinforced the relation in activity of all subregions of the hippocampus (DG, CA1 and CA3). Moreover, atVNS produced a marked enhancement in the correlated activity between hemispheres, especially in the frontal and amygdalar areas. A summary of intra and interhemispheric significant correlations for each condition is summarized in Table 1.

**Figure 3.**
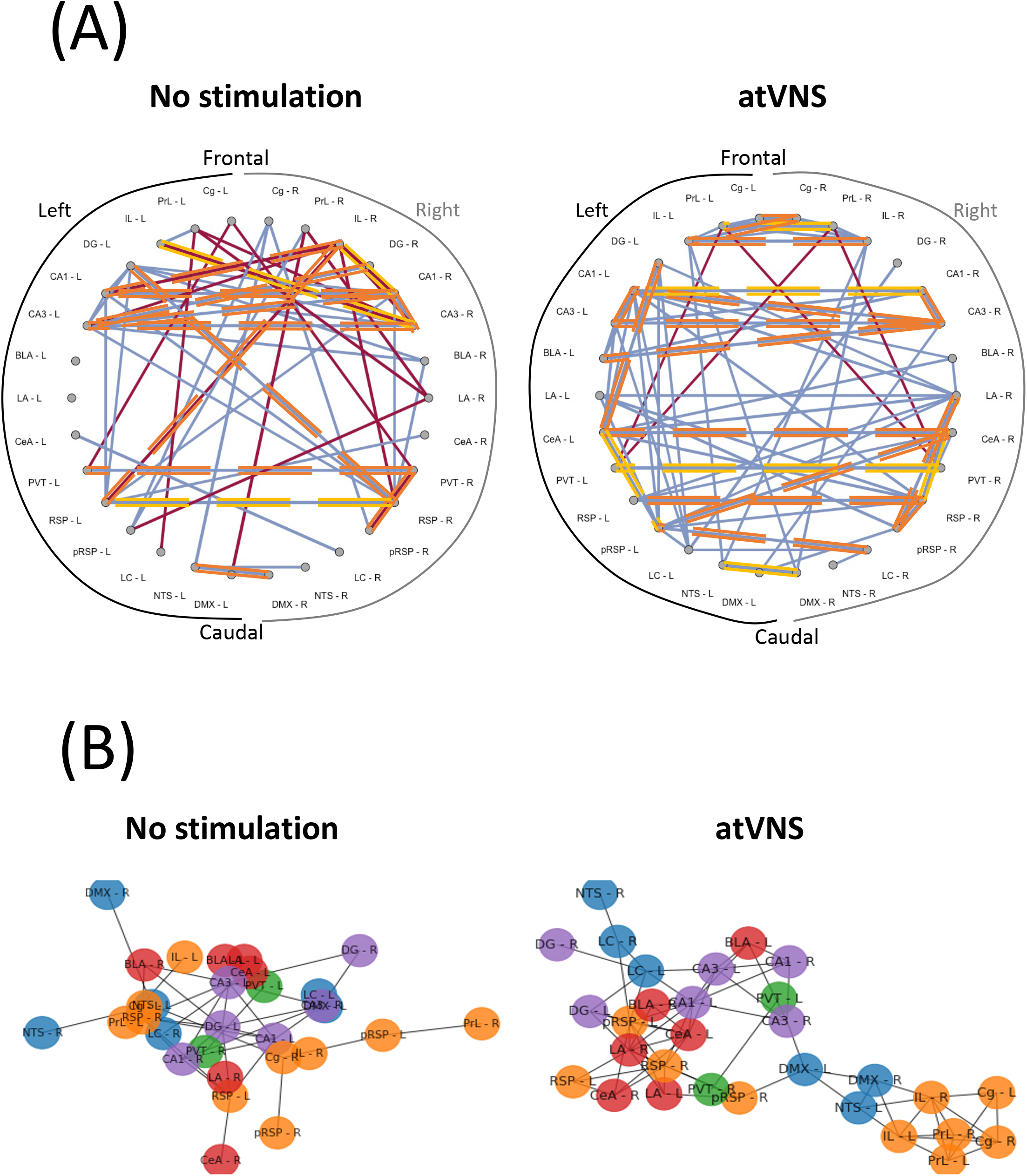
**(A)** Network connectivity graphs displaying only the significant correlations (p<0.05). Connecting lines represent Spearman correlation (positive correlation in blue, negative correlation in red). Strongest significant correlations are highlighted in orange (p<0.01) and yellow (p<0.001). Regions are presented from frontal to caudal and separating left and right sides. **(B)** Network connectivity Kamada-Kawai plots displaying only positive significant z-score for No stimulation and atVNS conditions. Colors represent cortical (orange), hippocampal (purple), amygdalar (red), thalamic (green) and brainstem (blue) groups. Regions are grouped based on the connectivity strength between them.

**Table 1.** Intra-hemisphere and inter-hemisphere number of significant correlations in No stimulation and atVNS conditions for frontal, hippocampal, amygdalar and brainstem areas.

| Areas | Intra-hemisphere |  |  |  | Inter-hemisphere |  |
| --- | --- | --- | --- | --- | --- | --- |
|  | No stimulation |  | atVNS |  | No stimulation | atVNS |
|  | Left | Right | Left | Right |  |  |
| Frontal cortex | 3 | 5 | 4 | 5 | 7 | 10 |
| Hippocampus | 5 | 3 | 11 | 1 | 10 | 8 |
| Amygdala | 0 | 1 | 5 | 4 | 5 | 11 |
| Brainstem | 1 | 0 | 6 | 2 | 3 | 5 |

Notably, while there was an overall increase in connectivity after atVNS, the prefrontal-retrosplenial axis, characteristic of the default mode network, was not observed in control conditions, and atVNS did not have any marked effects on engaging this axis (Supplementary Fig. 3).

Additionally, the network representation using Kamada-Kawai plots for No stimulation and atVNS conditions, revealed a relevant re-organization of the network due to atVNS with a more evident cross-talk between the brainstem areas and frontal and hippocampal regions (Fig. 3B). Indeed, atVNS produced a clear segregation of Cg, IL and PrL cortices with left NTS and DMX as connection nodes to the remaining structures. Also, the amygdaloid nuclei and the RSP cortex took a central role that was occupied by the hippocampal nuclei under No stimulation conditions.

Regarding the evaluation of global brain network metrics, a marked increase in all the evaluated properties was observed in atVNS brains (Table 2).

**Table 2.** Global network metrics for No stimulation and atVNS conditions. Global efficiency measures the proficiency of distant information transfer in a network; average clustering reflects the presence and prevalence of clustered connectivity, characteristics of segregated networks; average strength reflects the power of functional connections; average degree accounts for the amount of connections, related to network development and resilience.

|  | No stimulation | atVNS |
| --- | --- | --- |
| Global efficiency | 0.346 | 0.500 |
| Average clustering | 0.478 | 0.734 |
| Average strength | 4.037 | 5.956 |
| Average degree | 3.266 | 4.933 |

We further assessed the relative importance of the brain regions analyzed in the overall brain network based on regional network metrics. An increase in the relative importance of frontal areas in atVNS condition is observed, compared to No stimulation condition in which hippocampal regions showed a more relevant role. Furthermore, we found that brainstem regions, especially left LC, had a more relevant role in the atVNS network than in the No stimulation network, with a relative higher value of both nodal strength and nodal degree coefficients (Fig. 4A), since the brainstem regions constitute a passage between the vagus nerve afferents and superior brain regions. Indeed, if we compare the overall connectivity for LC region, we observe a significant increase in the connectivity of both left and right LC (Fig. 4B).

**Figure 4.**
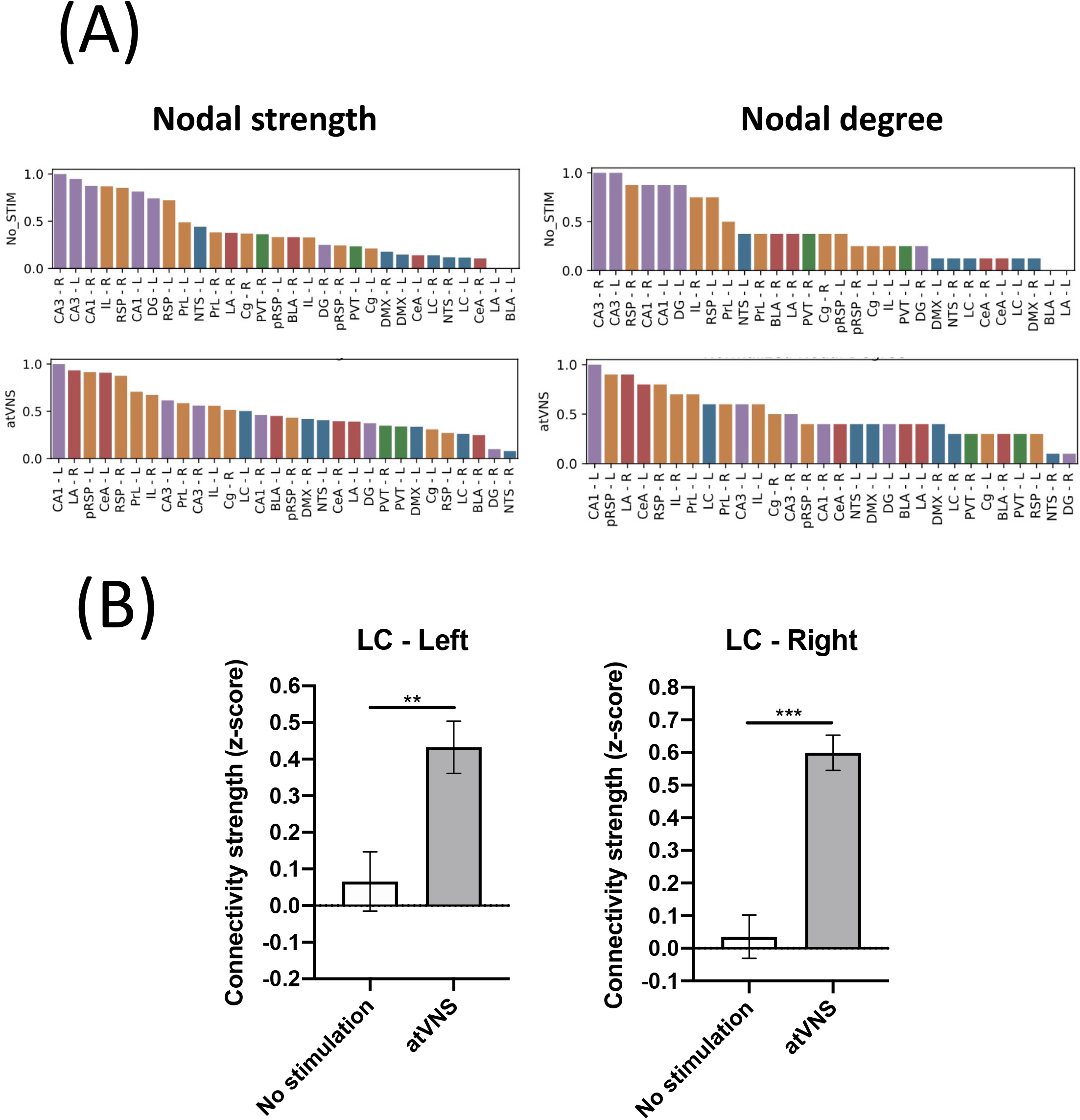
**(A)** Brain regions ranked in descending order based on nodal strength and nodal degree coefficients for No stimulation and atVNS conditions. Regions are displayed with color coding: cortex (orange), hippocampus (purple), amygdala (red), thalamus (green) and brainstem (blue). **(B)** Difference in the total connectivity of right and left *locus coeruleus* (LC) regions with the rest of the brain areas analyzed, presented as z-score and comparing No stimulation and atVNS conditions. ** p<0.01; *** p<0.001 by Kruskal-Wallis test.

## Discussion

Invasive and non-invasive vagus nerve stimulation (VNS) have been shown to modulate memory functions, both in animal [15,25,26] and human studies [27-29]. However, the presence of an effective time window for atVNS effectiveness, has not been explored before. Additionally, the brain activity outcome of atVNS under memory retention facilitated conditions is still under study.

Therefore, first we aimed to determine whether there was a critical time window for effective enhancement of object-recognition memory persistence through an acute session of atVNS. We found that atVNS in the *concha* of the left external ear of naïve CD-1 male mice, improved memory persistence, when the electrostimulation protocol was delivered immediately after the familiarization phase of the NORT. Conversely, atVNS delivered 3 h after the training of the NORT did not show any sign of effect on recognition memory at 48 h. This evidence points out the presence of an effective time window for atVNS efficacy in modulating memory persistence. This result is in agreement with a previous clinical report describing that the stimulation, in this case through an invasive approach, was effective when given around the learning phase of a Hopkins Verbal Learning Test, and not during the recall phase [28]. Therefore, both invasive and non-invasive forms of VNS, would principally enhance memory consolidation, leading to a better retention power when the electrical stimulus is delivered immediately after the familiarization or learning process.

Second, we investigated atVNS effects on neuronal and network activity, taking into consideration brainstem regions associated to vagal afferences, brain areas important for memory processes and brain regions implicated in the DMN. It is well established that memory is not stored in a single brain area, but in a network composed by multiple regions [30]. In the case of recognition memory, the hippocampal formation comprises the main brain region involved [31], although cortical and subcortical areas are also engaged [32]. It has been postulated that memories are initially retained in the hippocampus, and then the information is transferred into the neocortex where it can be consolidated and stored for longer periods [32,33]. Furthermore, the amygdala complex plays an important role in memory processes, especially under emotionally arousing experiences [34]. These brain regions are contacted by brainstem regions which are relevant for setting the stage concerning the responsiveness associated to the arousal state [35]. In this regard, the LC can convey the information from vagal afferents arriving to the NTS [7] and affect memory-related regions. Hence, we wondered whether the consolidation of new object-recognition memories, facilitated by atVNS, could be mediated by a change in neuronal activation or a re-distribution of the activity relation between brain areas.

Neuronal activation and functional connectivity were analyzed by computing the c-Fos density and interregional correlations across animals that received or not the electrostimulation procedure immediately after the familiarization phase of the NORT. We first calculated bilaterally the c-Fos density in a set of cortical, hippocampal, amygdalar, thalamic and brainstem regions for No stimulation and atVNS conditions. Notably, atVNS did not promote significant changes in c-Fos density in any of the thirty (fifteen per hemisphere) brain regions studied. Other previous reports found changes in c-Fos density after the electrostimulation, particularly an increase in the number of c-Fos+ cells, in areas of the brainstem [36-38]. However, all these studies used percutaneous or invasive VNS approaches, or applied the stimulation longer times. Our atVNS protocol instead is completely non-invasive, involves the auricular branch of the vagus nerve and the stimulation period is limited to 30 min, producing a significant effect in memory performance. Thus, the fact that no significant differences could be observed in the number of c-Fos+ cells in brain areas where other studies have found VNS-associated modulations, are probably due to the non-invasiveness and short stimulation protocol of atVNS procedure. Furthermore, several studies have been conducted using functional magnetic resonance imaging (fMRI) and atVNS, especially in humans, to shed some light on the possible mechanisms and brain networks involved during atVNS. In general, left atVNS produced a significant activation in the ipsilateral NTS, LC and prefrontal and cingulate cortices, while bilateral deactivation was found in the hippocampus and hypothalamus, and controversial results were described in the amygdala [39-42]. In contrast to fMRI procedure, we used c-Fos as a proxy for cellular activity; the main advantage of this approach is the cellular resolution mapping, although the poor temporal resolution is the principal limitation [43]. Notably, when we investigated c-Fos functional network, we found a significant reorganization, due to the electrostimulation procedure. atVNS resulted in an enhanced number of significant inter-hemisphere correlations compared to those observed in the No stimulation condition, especially in frontal and amygdalar areas. Moreover, significant correlations were found for sub-networks in the hippocampus (DG, CA1, CA3) and frontal areas (PrL, IL, Cg), with a specific increase in the correlation between the left LC and the dentate gyrus after atVNS. The distribution of the network differed between stimulation conditions. Under atVNS prefrontal cortices were segregated and linked to the remaining structures through DMX and left NTS, compared to No stimulation conditions, suggesting the ability of atVNS to dynamically reconfigure large-scale organization. In addition, a reorganization of the amygdala nuclei was observed pointing to a key role of this structure in the atVNS-mediated memory enhancement.

We also explored the effect of atVNS on selected DMN regions. This network has been also described in the mouse brain and involves the pRSP, RSP, Cg, PrL and IL areas as components of the network [13]. By studying these areas in both hemispheres separately we found that atVNS does not favor the connectivity of the DMN areas, as there is an absence of communication of the DMN anterior-posterior axis between frontal areas and retrosplenial cortex. This is in agreement with the idea that the DMN is mostly disengaged during task performance, and atVNS does not facilitate its engagement, and it may favor its disengagement.

Previous studies, using a combination of c-Fos expression and network analysis, found that long-term contextual fear memories are stored in a brain network composed by thalamic, hippocampal and cortical regions [24], while the brain network composed by hippocampus, medial prefrontal cortex, anterior cingulate cortex and amygdala was found required for the consolidation of social recognition memory [30]. These studies suggest that distinct types of memory are supported by exclusive functional memory networks that can be revealed by c-Fos analysis. Our findings in this study suggest that enhanced object-recognition memory consolidation is not prompted by an increase in neuronal activation by acute atVNS. Instead, we found a redistribution of the activity, and identified the correlation between brainstem nuclei and hippocampus and frontal areas as the privileged communication that may support the enhancement in memory persistence. Therefore, future studies should also focus on the real-time assessment of changes in neuronal activity induced by atVNS, to better understand its potential in modulating memory processes.

## Acknowledgements

We thank Dulce Real and Francisco Porrón for expert technical assistance and the Laboratory of Neuropharmacology-NeuroPhar for helpful discussion. The project that gave rise to these results received the support of a fellowship from “la Caixa” Foundation (ID 100010434). The fellowship code is LCF/BQ/IN18/11660012 (C.B.-P.). This project has received funding from the European Union’s Horizon 2020 research and innovation programme under the Marie Slodowska-Curie grant agreement No. 713673. This work was supported by the Ministerio de Economía, Innovación y Competitividad (MINECO) [#RTI2018-099282-B-I00 to A.O., #SAF2017-84060-R to R.M.]; the Instituto de Salud Carlos III [#RD16/0017/0020 to R.M.]; the Generalitat de Catalunya [2017SGR-669 to R.M.]; the ICREA (Institució Catalana de Recerca i Estudis Avançats) Academia to A.O., A.I. and R.M.; Grant “Unidad de Excelencia María de Maeztu”, funded by the MINECO [#MDM-2014-0370]; PLAN E (Plan Español para el Estímulo de la Economía y el Empleo). FEDER funding is also acknowledged.

## Author contributions

**C.B.-P**. participated in experimental design, conducted and analyzed experiments and wrote the manuscript. **E.M.-M**. produced network analysis and revised the manuscript. **I.G.-A**. conducted and analyzed experiments. **R.M**. participated in the supervision and experimental design, funded the project and revised the manuscript. **A.I**. participated in the supervision and stimulator design and generation, funded the project and revised the manuscript. **G.S**. participated in the supervision and analysis of network data, and revised the manuscript. **A.O**. conceptualized, participated in experimental design, supervised, funded the project and wrote the manuscript. All authors reviewed and approved the final version of the manuscript.

## Supplementary figure legends

**Supplementary Figure 1.**
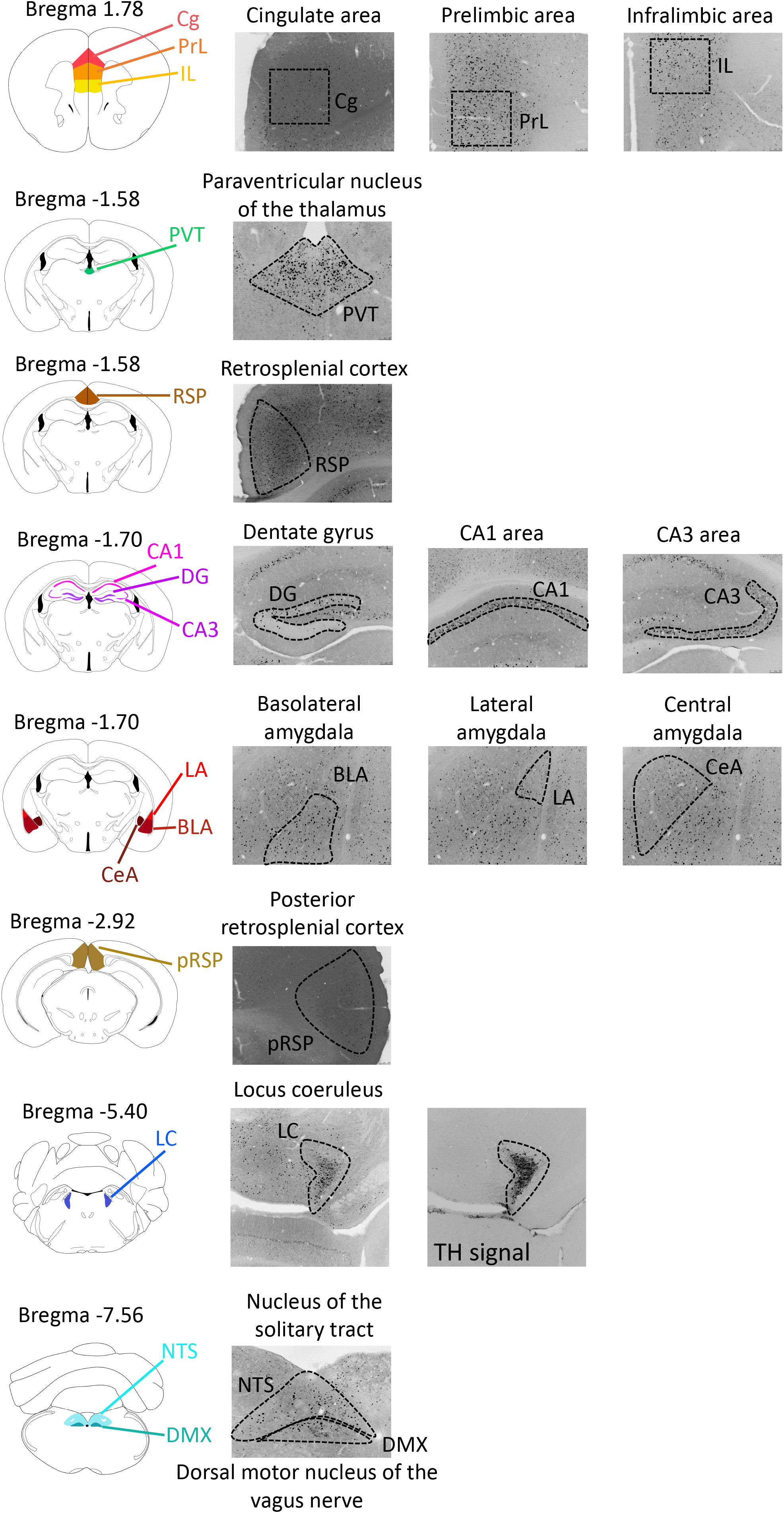
Schematic representation and representative pictures of c-Fos immunofluorescence of brain region analyzed. Brain areas are displayed from frontal to caudal with the corresponding coordinates relative to Bregma.

**Supplementary Figure 2.**
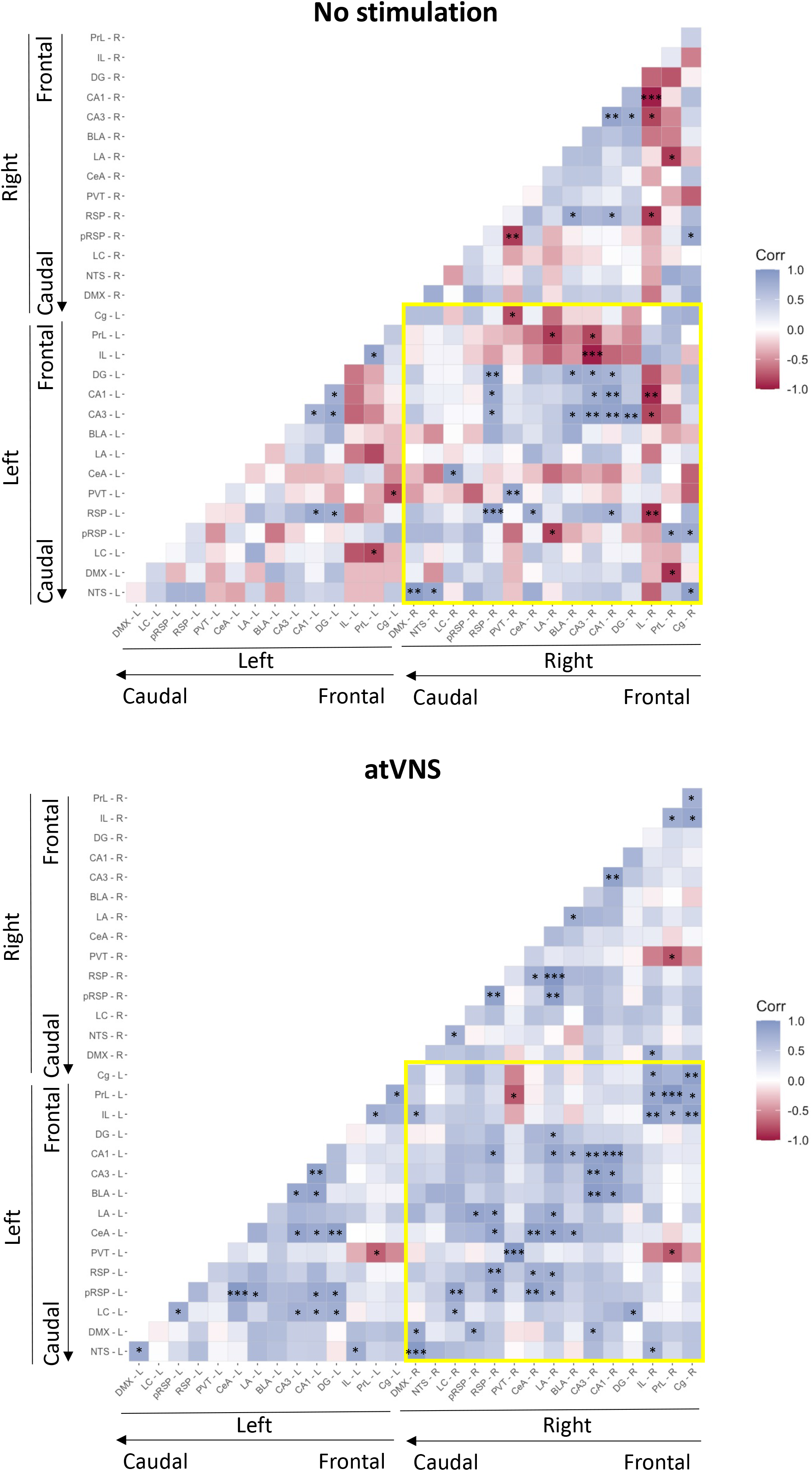
Connectivity matrices showing inter-regional Spearman correlations for c-Fos density. Axes represent brain regions organized from frontal to caudal and separating left and right sides. The yellow square denote the interhemispheric connections. Colors reflect Spearman correlation coefficients (scale above), significant correlation is marked by * p<0.05, ** p<0.01 and *** p<0.001.

**Supplementary Figure 3.**
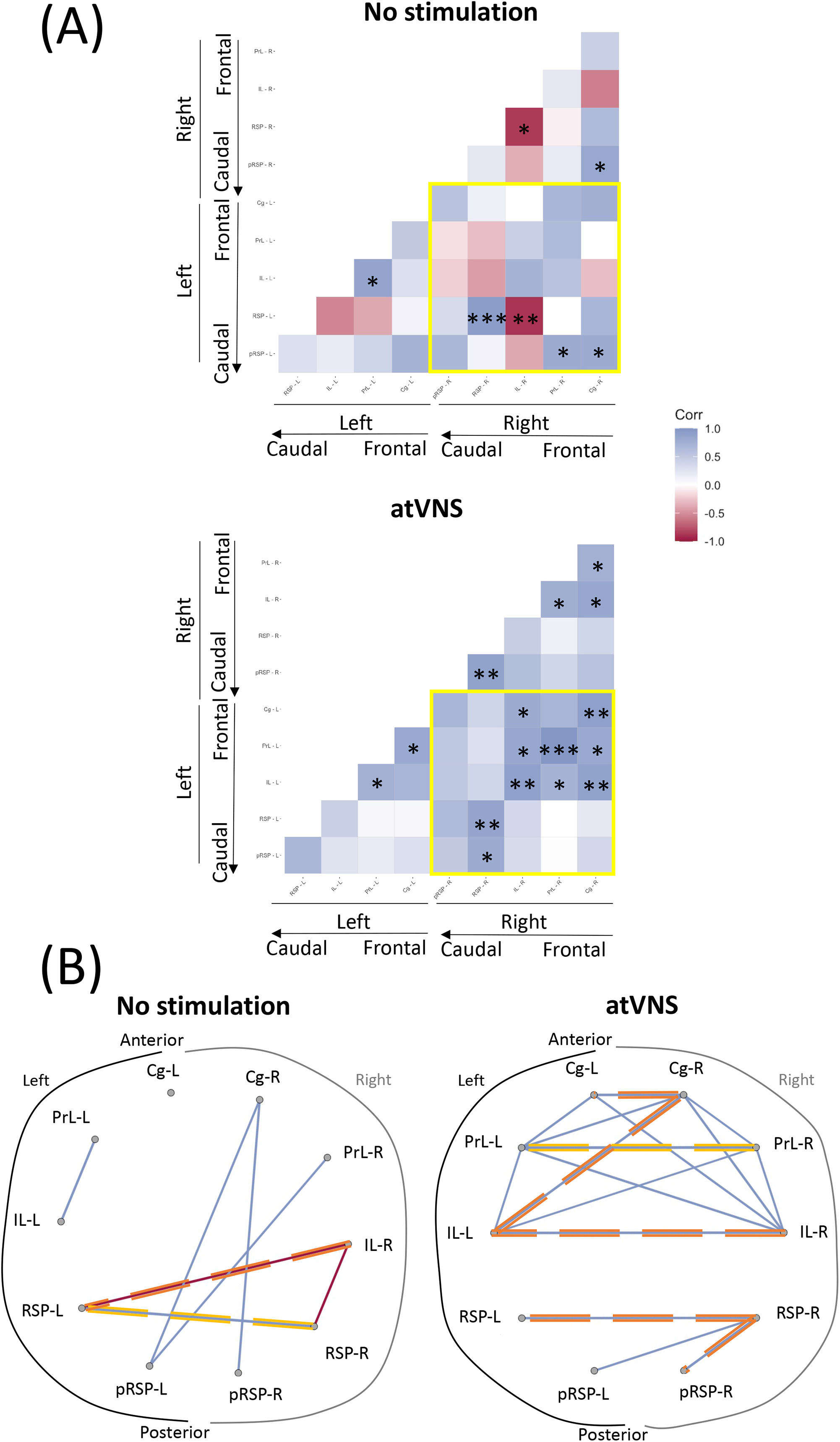
**(A)** Default mode network connectivity matrix. Axes represent brain regions organized from frontal to caudal and separating left and right sides. The yellow square denote the interhemispheric connections. Colors reflect Spearman correlation coefficients (scale above) and labels within squares correspond to p values of correlations (* p<0.05, ** p<0.01, *** p<0.001). **(B)** Circle plots showing significant correlations (p<0.05) in default mode network areas. Connecting lines represent Spearman correlation (positive correlation in blue, negative correlation in red). Strongest significant correlations are highlighted in orange (p<0.01) and yellow (p<0.001). Regions are presented from frontal to caudal and separating left and right sides.

